# 2.7 Å cryo-EM structure of vitrified *M. musculus* H-chain apoferritin from 200 keV “screening microscope”

**DOI:** 10.1101/738724

**Authors:** Farzad Hamdi, Christian Tüting, Dmitry A. Semchonok, Koen M. Visscher, Fotis L. Kyrilis, Annette Meister, Ioannis Skalidis, Lisa Schmidt, Christoph Parthier, Milton T. Stubbs, Panagiotis L. Kastritis

## Abstract

Here we present the structure of mouse H-chain apoferritin at 2.7 Å (FSC=0.143) solved by single particle cryogenic electron microscopy (cryo-EM) using a 200 kV device. Data were collected using a compact, two-lens illumination system with a constant power objective lens, without the use of energy filters or aberration correctors. Coulomb potential maps reveal clear densities for main chain carbonyl oxygens, residue side chains (including alternative conformations) and bound solvent molecules. We argue that the advantages offered by (a) the high electronic and mechanical stability of the microscope, (b) the high emission stability and low beam energy spread of the high brightness Field Emission Gun (x-FEG), (c) direct electron detection technology and (d) particle-based Contrast Transfer Function (CTF) refinement have contributed to achieving resolution close to the Rayleigh limit. Overall, we show that basic electron optical settings for automated cryo-electron microscopy imaging, widely thought of as a “screening cryo-microscope”, can be used to determine structures approaching atomic resolution.

**Highlights:**

- The 2.7 Å structure of mouse apoferritin was solved using a 200 keV screening cryo-microscope
- The apoferritin reconstruction was resolved without an energy filter, aberration correctors, or constant-power condenser lenses
- Comparison to available crystallographic and cryo-EM structures from high-end cryo-microscopes demonstrates consistency in resolved water molecules, metals and side chain orientations
- Although radiation damage is more prominent at 200 keV compared to 300 keV, this type of instrumentation is more accessible to research laboratories due to its compactness and simplicity

## Text

Single-particle cryogenic electron microscopy (cryo-EM) has revolutionized high-resolution structure determination of biomolecular assemblies [1]. The first high-resolution structure better than 3 Å was resolved in 2014 (EMD-6224) and communicated the following year [2]. The vast majority (94%) of protein complexes solved by cryo-EM since 2014 are at a resolution lower than 3 Å (Fig 1A); as of July 2019, only 275 high-resolution (better than 3.0 A) cryo-EM reconstructions have been deposited in the Electron Microscopy Data Bank (EMDB, https://www.ebi.ac.uk/pdbe/emdb/), for which 194 atomic models (70.5%) are available in the Protein Data Bank (PDB, https://www.rcsb.org/).

**Figure 1.**
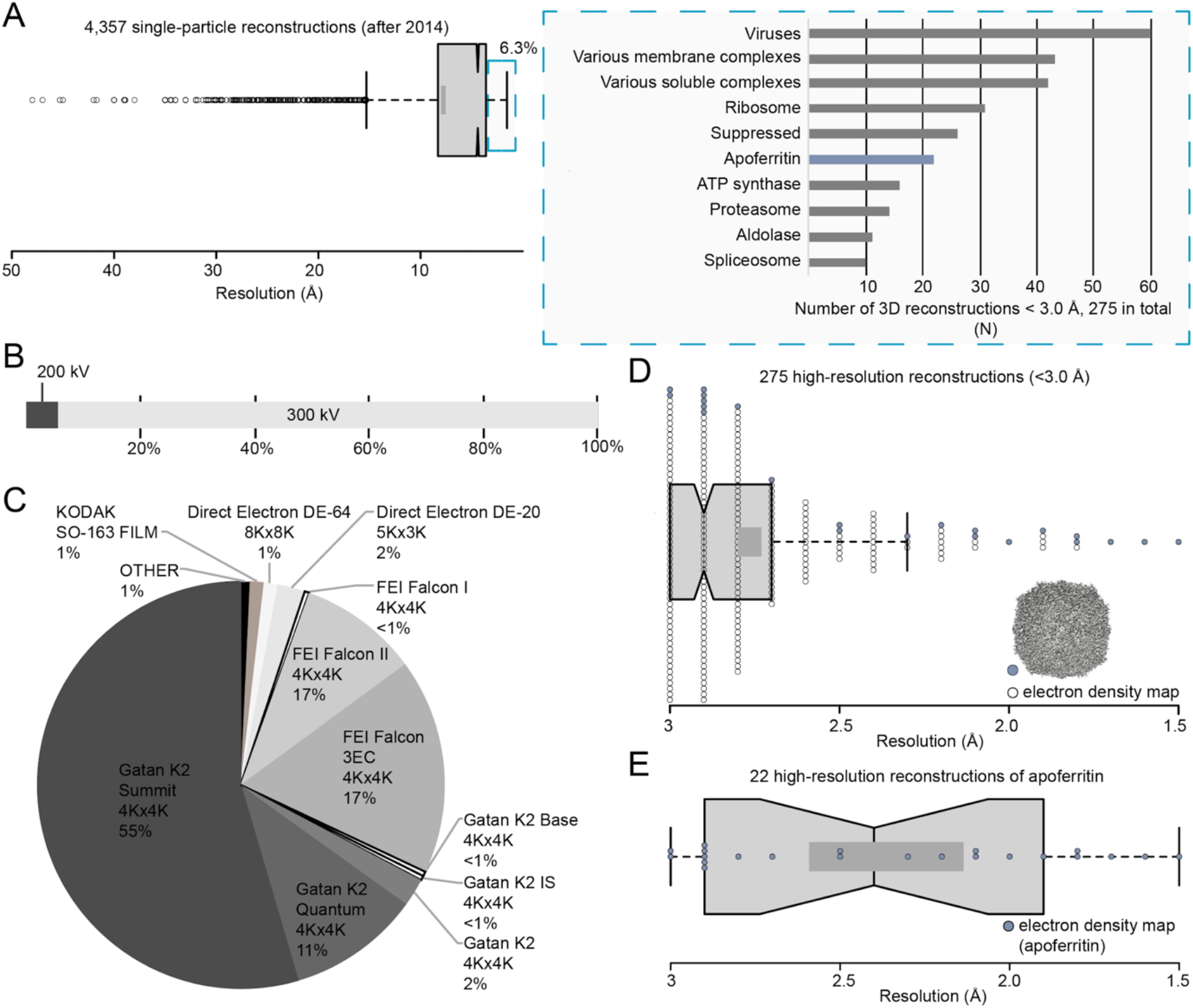
Analysis of single-particle cryo-EM structures from EMDB. (A) On the left, box plot shows the distribution of resolution in all single-particle reconstructions after 2014 and on the right, a bar plot shows the type of specimen reaching resolution better than 3.0 Å; (B) Percentage of structures reaching resolution better than 3.0 Å according to the electron microscope accelerating voltage used to acquire micrographs; (C) A pie chart showing the detector types applied for deriving molecular structures better than 3.0 Å; (D) A box plot showing the distribution of resolution for structures better than 3.0 Å; apoferritin reconstructions are highlighted in blue; (E) A box plot showing the distribution of resolution for apoferritin reconstructions. Dark grey rectangles in (D) and (E) show the confidence intervals at 83%; middle black line shows the average. Data points are shown with circles.

Statistical analyses on the 275 reconstructions demonstrate a preference towards specific biomolecules systematically reaching high resolution, with 60% relating to 7 types of molecules (Fig 1A). 164 of these reconstructions are symmetric (Fig S1), of which half are isometric (59 icosahedral, 25 octahedral and 2 tetrahedral). The overwhelming majority of high-resolution reconstructions (95%) have been resolved using high-end, automated 300 keV electron microscopes (Fig 1B), equipped with direct electron detectors (98%) (Fig 1C), energy filters and constant-power condenser electromagnetic lenses. The high frame rates of direct detection devices (DDD) allow compensation for electron beam induced particle motion [3], energy filters contribute to image enhancement by removing inelastically scattered electrons that contribute to background noise [4], while constant-power condenser lenses allow the switches to be made in optical settings necessary during high-resolution low-dose imaging protocols without affecting beam stability. In combination with advanced image processing algorithms [5], [6], [7], [8], such a highly sophisticated set-up is capable of achieving resolutions between 3.0 Å to 2.5 Å, with a few reconstructions surpassing 1.8 Å resolution. Interestingly, most of the reconstructions resolved at a resolution of 2 Å or better correspond to apoferritin from various organisms (Fig 1D, Fig 1E, Table S1), as a result of its high symmetry and intrinsic stability. Indeed, apoferritin is now commonly used as a standard sample to assess microscope performance and implement developments in cryo-EM [5], [9], [10], [11].

High resolution cryo-EM studies are therefore most likely to succeed for stable specimens imaged using high kV electron microscopes with energy filters and direct detection technology, in combination with modern image processing routines [12]. Such sophisticated equipment is however expensive, requiring high-level strategies to maintain microscope stability and performance, so that the method would appear to be accessible in a daily manner to only few laboratories. Recent publications have explored the potential of 200 kV microscope to obtain reconstructions higher that 3 Å [13], [14] [15], [16] and [17], see Table S2. The dedicated instrument used to resolve those molecular structures is, again, highly sophisticated, and is in addition, costly. The set-up requires dedicated space due to its size and is equipped with advanced direct electron detection technology (DDD) with high frame rates. This DDD has been shown to resolve more than half (55%) of high-resolution electron density maps (Fig 1C).

Below, we describe the differences between the instruments above and our electron optical settings. We show that basic settings for automated cryo-electron microscopy imaging, corresponding to what is widely considered as a “screening cryo-microscope”, can be used to determine structures at high resolution. We use mouse H-chain apoferritin to produce an atomic model resolved at 2.7 Å (see Supplementary material for methodological details) and compare it to the corresponding 2.24 Å crystal structure (PDB ID: 3WNW [18]).

Why are 200 keV microscopes generally considered to be sub-optimal for high resolution studies? Apart from the reduction in the high angle diffraction resolution limit at lower energy [19], the probability of electrons being inelastically scattered increases with decreased keV. This leads to an increase in the background noise of the acquired micrographs, as well as increased interactions with the sample. This in turn increases the likelihood of electron-induced radiation damage [20], [21], although inelastic scattering can be compensated for by the use of an energy filter, as demonstrated recently for mouse apoferritin [22]. On the other hand, 200 keV instruments offer distinct advantages for cryo-EM applications, including the higher contrast of the specimen [21], [23] and considerably lower maintenance requirements due to the simpler design.

The electron microscope used here is equipped with an automated loading mechanism that minimizes manual intervention during the loading procedure and hence ice contamination. The illumination system includes a stable high-brightness Field Emission Source (x-FEG) and two basic condenser electromagnetic lenses, which lowers the cost of the system. Care must be taken to adjust and maintain beam parallelism, however. The objective lens is a constant power symmetric twin lens equipped with an auxiliary lens (the micro-condenser lens), which provides a narrow parallel beam for localization of the emission in the field of interest while avoiding any damage to the surrounding area. The microscope column is closer to the ground compared to other 200 keV and 300 keV microscopes, providing increased mechanical stability and reduced sensitivity to external vibrations (although complicating fitting of either an in-column or post-column energy filter). The more compact system allows installation in standard laboratory rooms, resulting in increased accessibility and reduced infrastructure costs to control environmental conditions (e.g. field cancellation device, humidity, temperature, noise etc.) that are critical for microscope stability [24]. Otherwise, standard practices were followed in the microscopy site.

A DDD in counting mode for used for imaging, with 40 frames per second (fps) of 4Kx4K and an electron dose of 0.97 e-Å^−2.^s^−1^. Vitrification at high concentration (4 mg.ml^−1^) allowed imaging of a suitable number of particles (Fig S2A) on only 300 micrographs in less than 12 hours. Frames were motion corrected with built-in microscope software frame alignment routines followed by frame alignment procedures in Relion 3.0.5 [5]. Final aligned stacks show global motions ~0.3 Å on average (Fig S2A). Applied and calculated defocus values varied between 600 nm and 1,600 nm and showed high consistency when calculated with gctf [25] (Fig S2B), demonstrating the robustness of the defocus calculation. Defocus values close to the Scherzer defocus allowed us to perform contrast transfer function (CTF) fitting with high confidence using gctf [25] (Fig S2C). This resulted in consistent recording of electron micrographs with calculated average resolution of ~3.2 Å (Fig S2B).

Standard image processing procedures were performed in Relion 3.0.5 (Fig 2A-E, Fig S2) to calculate an electron optical density reconstruction of mouse apoferritin at 2.7 Å (Fig 2E). In particular, iterations of particle-based CTF refinement, followed by particle-based polishing and beam-induced motion correction procedures [26] were critical factors in increasing the resolution of the final reconstruction of mouse apoferritin from 3.6 Å to 2.7 Å (Fig 2E).

**Figure 2.**
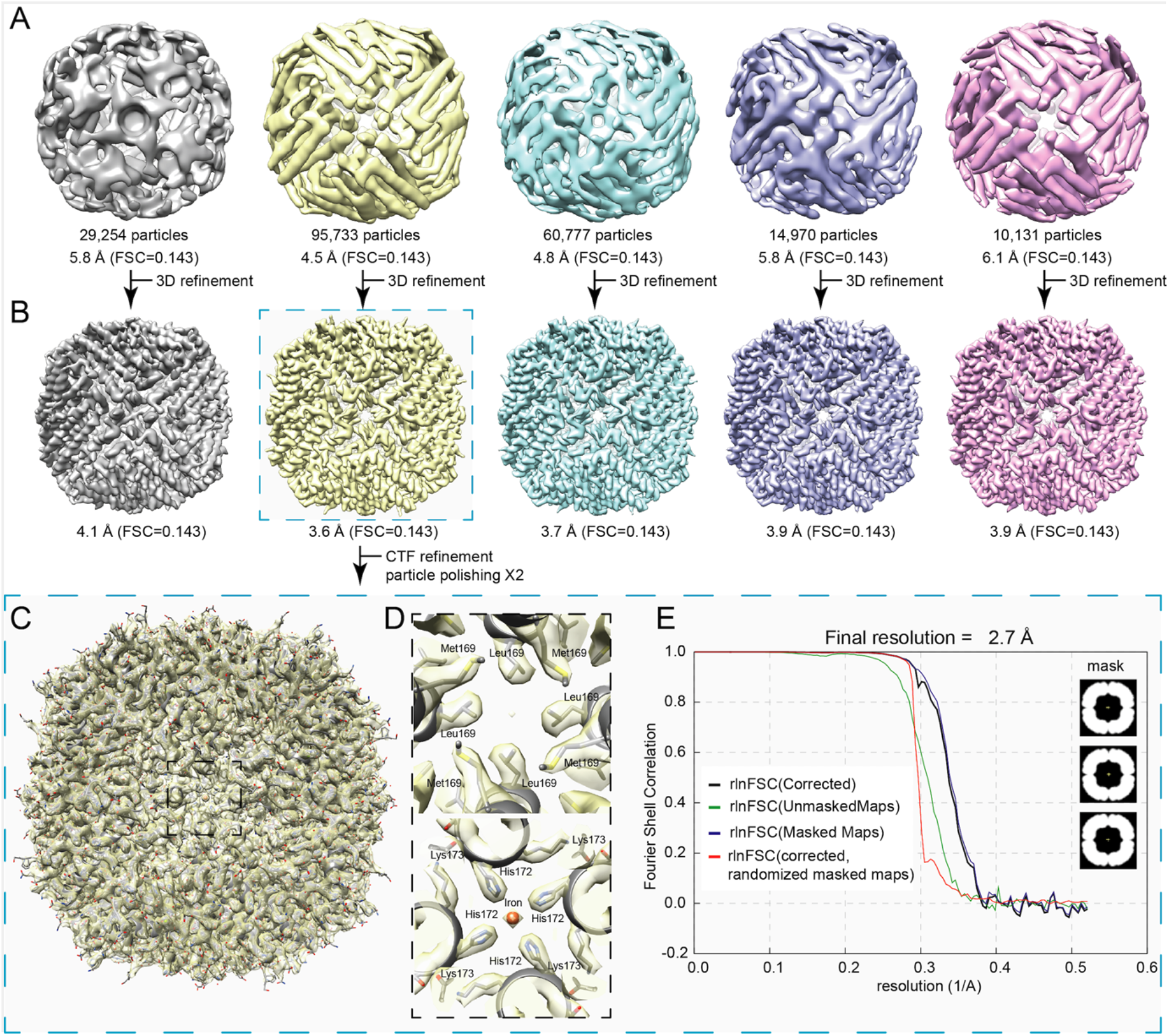
Image processing of single-particle data after 2D classification and final resolution estimation. (A) Five (5) 3D classes were initially calculated, reaching resolutions from 4.5 Å to 6.1 Å. (B) Subsequent 3D refinement procedures led to reconstructions of the classes in a resolution range of 3.6 Å to 4.1 Å. (C) Overlap of the final atomic model of mouse apoferritin with the refined Coulomb potential density map. (D) Close-up images along the 4-fold axis of apoferritin at different depths of the protein shell. Coulomb potential densities are recapitulated for the corresponding channel, including a bound iron atom. (E) Fourier shell correlation plot for the final 3D reconstruction shown in (C). At an FSC of a reported correlation value of 0.143, resolution reaches 2.7 Å.

Overall, the architecture of mouse apoferritin is as expected (Fig 2C), and two layers along the 4-fold channel axis in which an iron atom is bound is illustrated along with the resolved densities in Fig 2D. The Coulomb potential map clearly resolves densities for all secondary structure elements of apoferritin (helices A-E and Loop L (Fig 3)), with the appearance of main chain carbonyl oxygens consistent with the FSC-calculated resolution. Moreover, side chain densities are clearly resolved, with the exception of acidic side chain carboxylates. Nevertheless, densities for the latter can be observed at lower contour levels, *e.g*. for Asp131 and Glu134 (Fig 3 and Fig S2). The reduced density may be due to the enhanced sensitivity of acidic side chains to electron radiation and subsequent damage [27], [28], which is more prominent at 200 keV compared to apoferritin structures resolved at 300 keV [5], [9]. Other residues, such as polar (Asn98, Gln112) and aromatic (Trp93) amino acids are well resolved independent of contour level. The quality of the reconstruction is underlined by the unambiguous observation of *cis*Pro161 (Fig S3).

**Figure 3.**
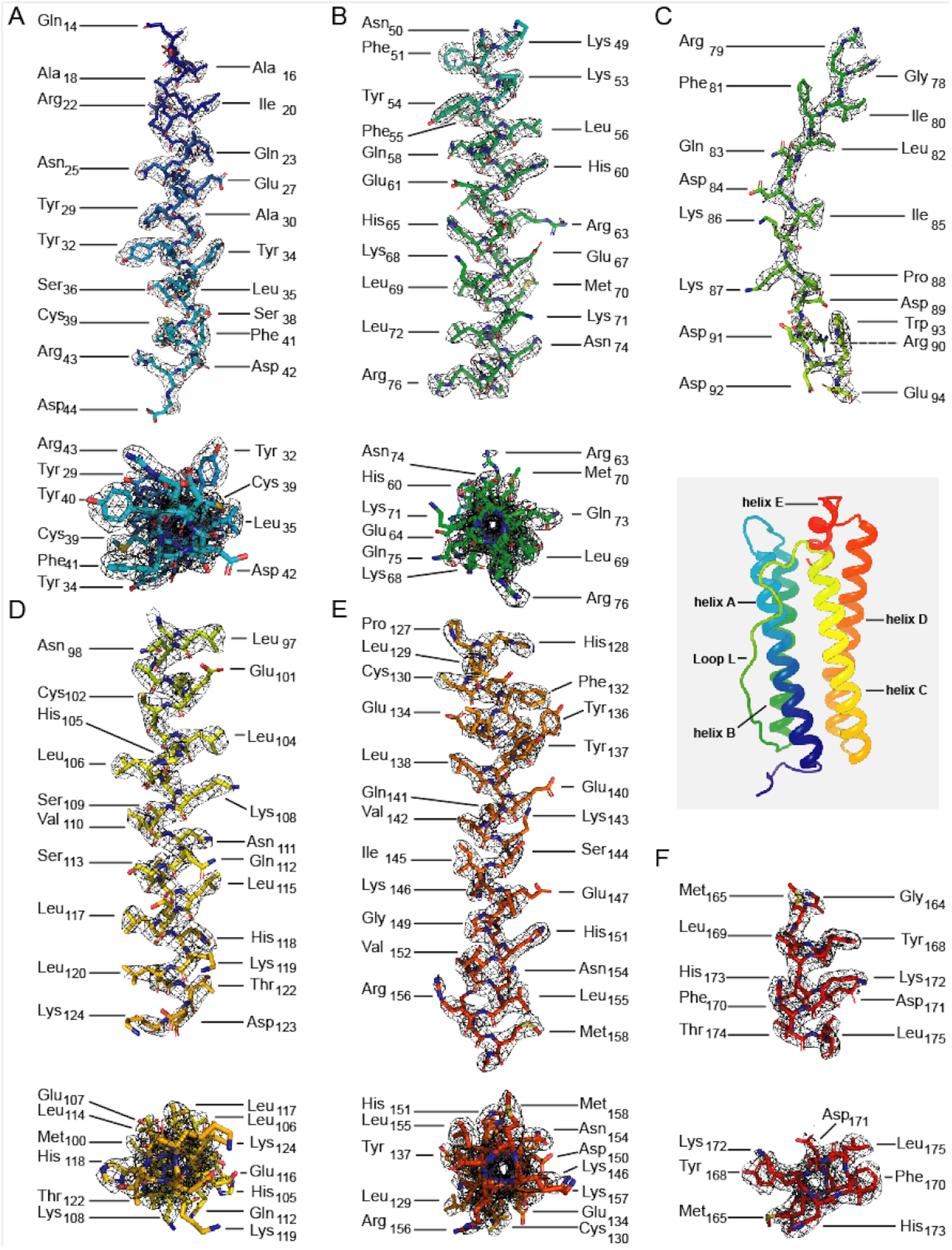
Coulomb potential densities for each secondary structure element of the apoferritin monomer (insert), with individual amino acid residues shown in stick representation. (A, B, D-F) helices A-E are shown with the fitted model and the corresponding density perpendicular to (top panels) and along (bottom panels) the helix axes. (C) Density and fitted model for loop L.

The crystallographic structure of mouse apoferritin has been solved with 12 independent monomers in the asymmetric unit (PDB ID 3WNW). Overall, our monomeric (octahedrally averaged) structure of H-ferritin shows only marginal differences to the individual monomers of the X-ray crystal structure (Fig 4A-D) with very low overall RMSD values (Fig 4A). Significantly, multiple side chain conformations could be distinguished in the map consistent with the crystal structure, e.g. Arg63 (Fig 4D) and Ile133 (Fig. 4D). While the structures display the same overall main chain conformation, side chain rotamers of a few amino acids (in particular charged side chains) display a broader distribution (Fig 4A). This is likely due to their predominant surface location, leading (in both crystal and cryo-EM structures), to statistical / dynamic fluctuations in solvent exposed regions [29], although electron induced radiation damage may also play a role.

**Figure 4.**
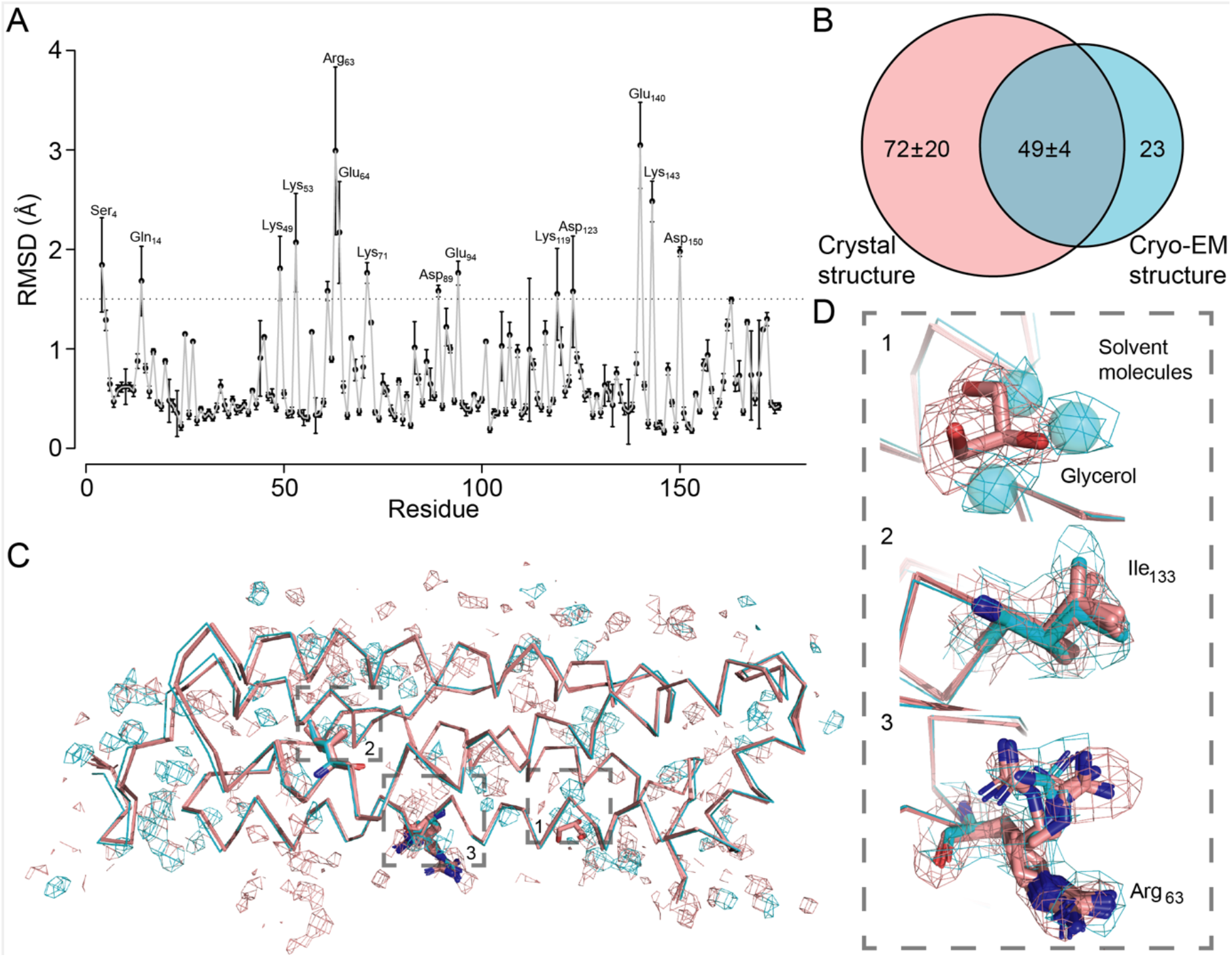
Comparison of the cryo-EM-derived structure to its crystallographic counterpart. Note that there are 12 monomers in the crystallographic asymmetric unit, so that comparisons are shown against all. (A) per-residue side-chain root-mean-square deviation (RMSD, Å) for all atoms. Residues exhibiting RMSD values >1.5 Å are highlighted in the plot. (B) Venn diagram showing overlap of solvent molecules derived from the cryo-EM map with those of the crystallographic monomers. (C) overlay of Ca atoms from the cryo-EM model (cyan) and the crystallographic monomers (pink) together with corresponding densities for the solvent. Boxes 1-3 denote positions shown in (D). (D) Comparison of cryo-EM (cyan) and X-ray (pink) models for selected residues with corresponding density/densities.

72 solvent molecules could be identified in the cryoEM map, compared to an average of 121 +/− 24 water molecules per monomer of the asymmetric unit in the X-ray structure (Fig 4B). Of these, 49 +/− 4 (68.1%) of the positions overlap, demonstrating the congruence of the cryo-EM and X-ray crystallographic solvent molecules, pointing to a structural role for these solvent molecules. For the remainder, some of the differences can be attributed to displacement of bound water molecules by glycerol (used as cryoprotectant) in the crystal structure (Fig 4D).

The biological function of apoferritin is to bind and store iron by combining with ferric hydroxide-phosphate compound to form ferritin. Various channels have been identified in structures of apoferritin that support this function, at the 4-fold, 3-fold and 2-fold axes [30],[31]. Comparison of the resolved channels in the apoferritin structure with those of their crystallographic counterpart shows that they are highly similar, clearly resolved in the cryo-EM model, with corresponding amino acid side chain conformations.

Application of our imaging protocols to mouse apoferritin demonstrates that our cryo-EM model is in agreement with its X-ray crystallographic counterpart. The overall fold, main chain organization, side chain and solvent molecule densities are readily interpretable from the reconstruction. In particular, functional channels of the molecule are well resolved. Nevertheless, the densities for negatively charged amino acid residues are consistently lower in the cryo-EM map, a phenomenon that is likely to be more prominent in 200 keV than in 300 keV reconstructions due to increased inelastic scattering. Another option could be that, due to the negative charge of the oxygen which may result in attenuated positive potentials, its corresponding density is simply not seen [32–36]. Our observations are consistent with previously resolved cryo-EM maps derived from 200 keV electron microscopes [13], [14]. We expect that use of an energy filter should allow achievement of higher resolution, as demonstrated for the recent reconstruction of mouse apoferritin at 2.0 Å, where images were recorded with a higher frame rate detector and a more sophisticated 200 keV microscope [22].

Overall, we expect that similar resolution can be achieved using the “screening microscope” on the condition that the sample and the vitrification process are optimal. Naturally, this resolution will be harder to achieve for samples with increased heterogeneity or ice thickness. Nevertheless, we have demonstrated that our electron optical settings and image processing methods allow structure determination of vitrified biological macromolecules at 2.7 Å using a “standard screening microscope”, providing an affordable option for in-house high-resolution structural biology.

## Accession numbers

The coordinates of the *M. musculus* apoferritin model are available from the Protein Data Bank (entry 6SHT), and the corresponding EM map is deposited in the Electron Microscopy Data Bank (EMDB entry EMD-10205).

## Acknowledgements

This work was supported by the Federal Ministry for Education and Research (BMBF, ZIK program) [grant number 02Z22HN23 to P.L.K.]; the European Regional Development Funds for Saxony-Anhalt [grant number EFRE: ZS/2016/04/78115 to P.L.K. and M.T.S.] and the Martin Luther University Halle-Wittenberg. The authors acknowledge the Thermo Fisher engineers Hartmut Lepuschitz, Christian Heiden, Volker Irmer and Dr. Sonja Welsch for installation of the cryo-microscope and expert technical assistance. The authors also thank engineer Mr. Mohammad Daraei for room preparations and field cancellation installation, Mr. Jürgen Simon and Mr. Ingo Schneider of the MLU as well as Prof. Dr. Reinhardt Paschke, Mr. Matthias Harm and Mr. Grossman of the Biozentrum for the necessary room modifications as well as supervision of the microscope installation, and our IT department, in particular Mr. Patrice Peterson, for excellent maintenance of the IANVS cluster.

