## Supplementary for "2.7 Å cryo-EM structure of vitrified *M. musculus* H-chain apoferritin from 200 keV “screening microscope”"

#### Supplementary Materials and Methods

Sample preparation and grid screening. A 3µl aliquot sample of 4.0 mg/ml concentration of mouse H-chain apoferritin was kindly provided by Thermo Fisher Scientific. The sample was applied to glow-discharged Quantifoil holey carbon grids (R1.2/1.3, 200 mesh). Cryo-EM grids were prepared using Vitrobot Mark IV System (Thermo Fisher Scientific) with 95% humidity, ashless filter paper (Standard Vitrobot Filter Paper, Ø55/20mm, Grade 595) and blotting time of 4 sec.

Cryo-EM data collection. Grids were imaged using a 200 keV Thermo Scientific Glacios Cryo-Transmission Electron Microscope (focal length 3.4 mm, objective aperture 100 µm) equipped with an XFEG source. Humidity and temperature of the microscope environment were controlled with a regular, commercially available air-conditioning system, and DC and AC magnetic fields of the room were actively cancelled with a dedicated magnetic field cancellation system. Images were recorded using a Falcon 3EC direct electron detector in counting mode. The images were collected using EPU software with an under focus range from 0.6 µm to 1.6 µm.

Microscope settings. Standard illumination alignments were performed prior to data collection. The gun was centred and tilted to have a central on-axis beam of electrons. The extracting voltage was set to the minimum of 4.2 keV to achieve a concentrated beam of low energy spread [1]. The strength of the first condenser lens (C1) was adjusted to achieve the desired dose rate of 0.93 electrons per pixel per second (e-/pix/sec). The second condenser lens (C2) was used for parallel beam illumination, achieved by optimizing the sharpness of the objective aperture and simultaneously adjusting the intensity of the electron beam to the highest possible illumination in diffraction mode. The sample for this adjustment was amorphous carbon areas of the

Quantifoil grid, imaged at eucentric height and defocus of -1 µm. The C2 aperture was set to 50 µm as beam defining aperture to minimize the beam diameter corresponding to the field of view, yielding a beam diameter of ~1.7 µm. Astigmatism of the C2 lens was corrected. The objective lens complex was also precisely aligned. A 100 µm aperture was symmetrically set in the focal plane of the objective lens for limiting high-angle inelastic or multiple scattered electrons. In this configuration, the half-beam collecting angle of the objective lens (β) was 14.7 mRad, giving a Rayleigh resolution criterion (maximum resolution cut-off) of:

$$r = \frac{0.61\lambda}{\beta} = 2.08\ \text{Å} \qquad (1)$$

where λ is the electron beam wavelength obtained from the relativistic de Broglie equation. Note that, although widely accepted, equation (1) is considered to be an underestimate in the electron microscopy literature [2]. Objective astigmatism and coma were minimised iteratively using the autostigmate and autocoma routines in EPU on the amorphous carbon region of the grid. 300 4KX4K movies were collected with a Falcon 3EC direct electron detector in counting mode, using EPU 2.1 for automated data collection. The pixel size was set to 0.96 Å. Exposure was set to 0.93 e-/Å$^2$/sec and 30 frames were collected in total, with an overall dose of 28 e-/Å$^2$. Applied defocus varied from -0.8 µm to -1.6 µm. Monitoring of image properties, estimated defocus and resolution was performed on-the-fly with the WARP software (https://www.biorxiv.org/content/10.1101/338558v1).

Image processing and map calculation. Movies were imported and analyzed in Relion 3.0.5. and motion correction was performed by default. To calculate the CTF, gctf was employed with standard parameters. We initially used 1 CTF-corrected micrograph to manually pick 113 particles with the box size set to 256 pixels. Iterative 2D classification was performed with an applied circular mask of 200 Å. In the first iteration of 2D, using the 113 particles, the most populated class was selected for template-based particle picking, low-pass filtered to 20 Å. In a test set of 7 micrographs, 5,059 particles were picked; visual inspection showed that >80% of particles were captured per micrograph. A second iteration of 2D classification was performed. The top 2 classes including 76% of the particles were again selected for template-based particle picking using default parameters. In all, 211,177 particles were auto-picked from 300 micrographs, and a third final iteration of 2D classification discarded 312 single particles, resulting in 210,865 individual particles. 3D classification in 3 classes was performed with an applied circular mask of 140 Å. We used as reference the 1.65 Å human apoferritin map resolved with Relion 3.0 (EMDB 0144), low-pass filtered to 30 Å. Octahedral O symmetry was applied. Resulting classes reached < 6.1 Å resolution, with the best and most populated class reaching 4.5 Å (45% of particles). Subsequent refinement of the best class reached an initial resolution of 3.6 Å. After 2 iterations of particle-based CTF refinement and particle polishing implemented in Relion 3.0, the final reconstruction of mouse apoferritin included 95,733 single particles and reached 2.73 Å resolution (FSC=0.143).

Model building and refinement. One protein monomer from the tetragonal crystal form of mouse H-chain apoferritin (PDB: 3WNW [3]) was placed into the octahedrally averaged cryo-EM Coulomb potential map $\rho_{map}$ as a rigid body using Chimera [4] and the polypeptide chain was fitted in real space using Coot [5].

#### Supplementary Figures

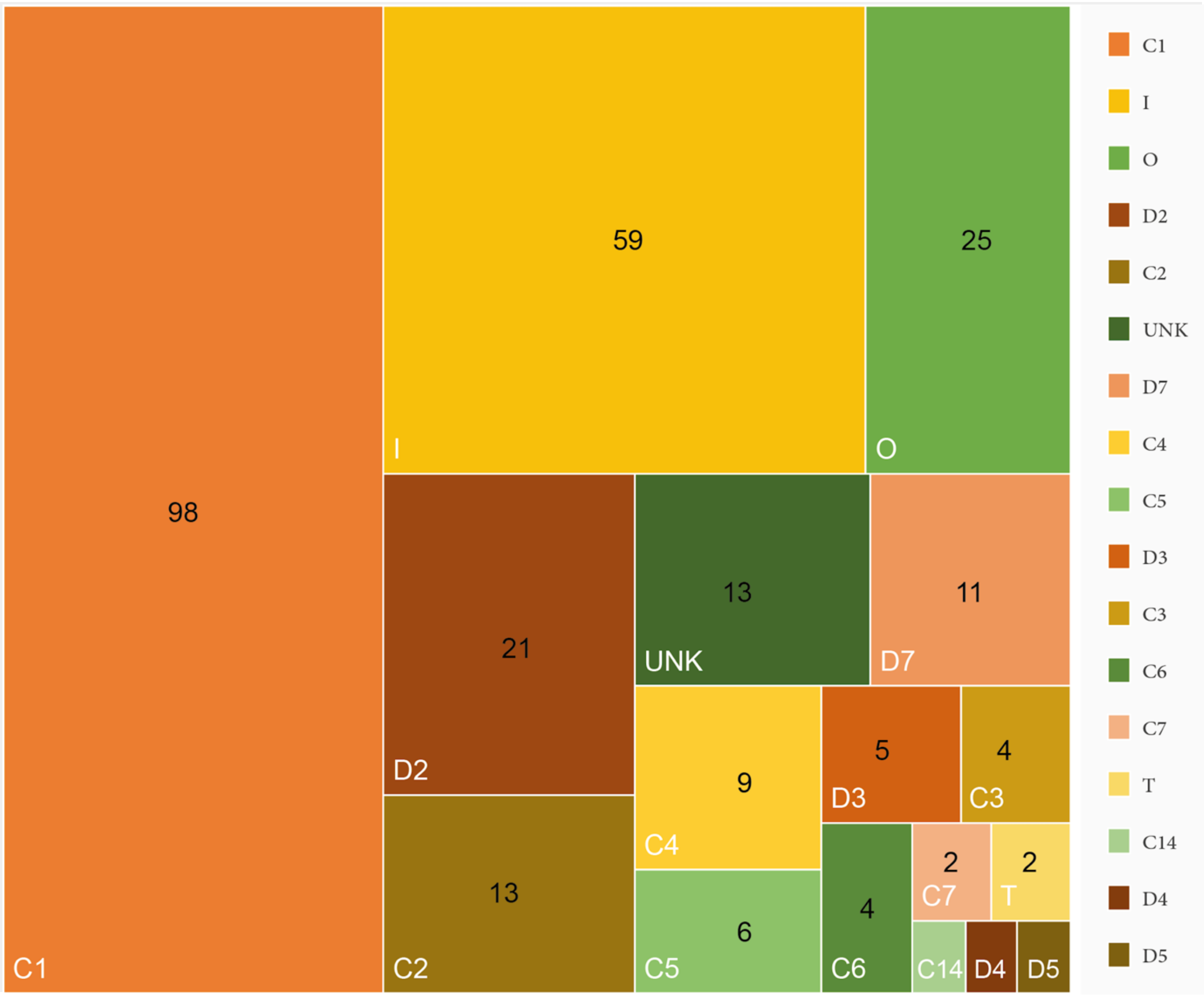


Figure S1. Matrix highlighting the statistics of point group symmetries of cryo-EM maps solved at resolutions better than 3.0 Å. The whole square shows the 275 structures, and the size of the different boxes highlights the structures resolved with the corresponding symmetry; each symmetry type is indicated with its corresponding color, as shown in both the matrix and the insert. 31 of the C1 (asymmetric) reconstructions correspond to those of ribosomes.

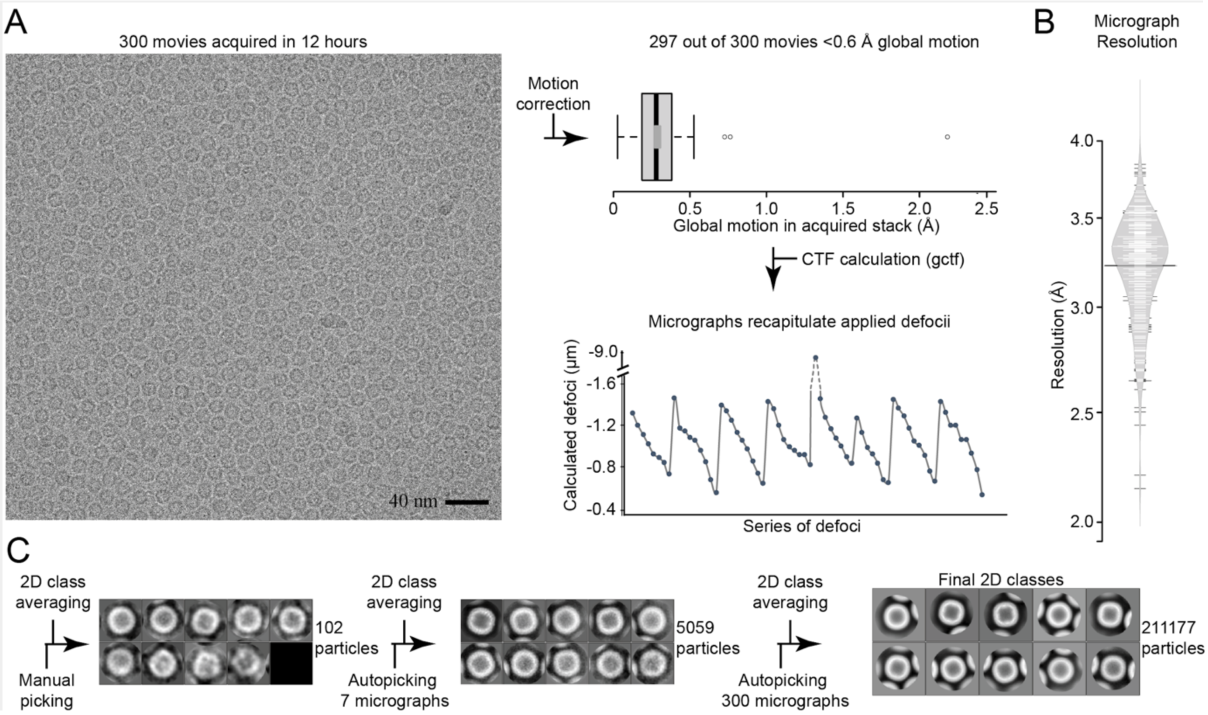


Figure S2. Statistics of acquired micrographs and initial 2D classification of apoferritin particles. (A) Left: a typical cryo-EM micrograph with recognizable apoferritin particles at a concentration of 4.0 mg/ml. Right: double-corrected motion corrected values are shown for the 300 acquired images as a bar plot. Bottom right: a series of calculated defocus values for the acquired micrographs is shown that recapitulate the applied defocus values determined for the data acquisition. (B) A bean plot showing the distribution of calculated micrograph resolutions; the black line shows the median; white lines represent individual data points; grey polygons represent the estimated density of the data. (C) 2D classification procedure to collect single-particles from all micrographs and subsequently select the 211,177 particles that were classified in five (5) classes in 3D (Fig. 2).

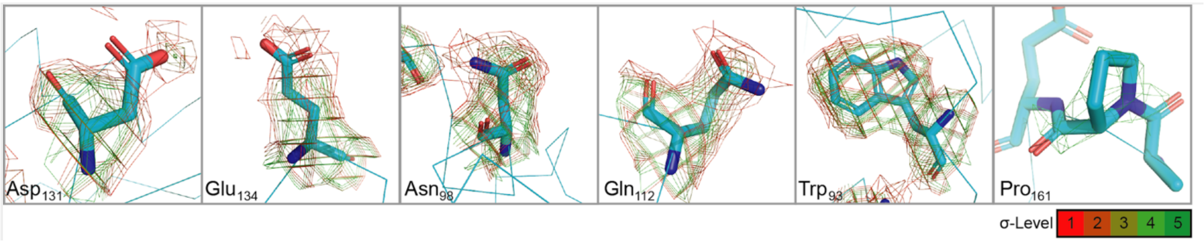


Figure S3. Side chains of helix E at different contour levels of the EM density (see text for details).

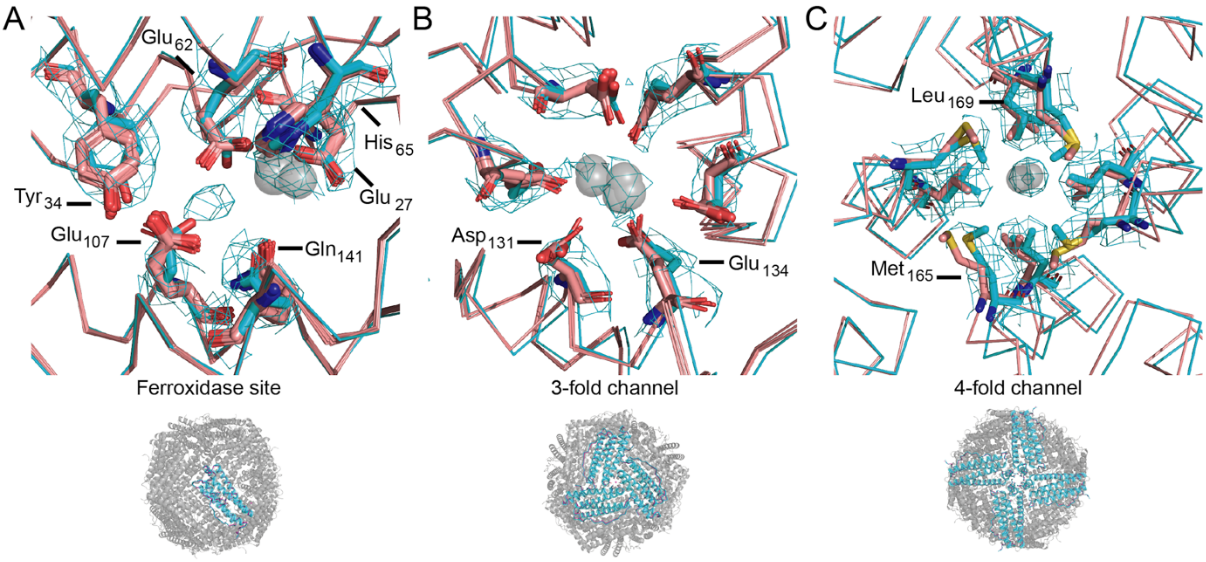


Figure S4. Apoferritin metal ion (grey transparent spheres) binding sites from the cryo-EM reconstruction (cyan) overlaid with corresponding residues and metal ions observed in the crystal structure (pink). (A) ferroxidase site ($Mg^{2+}$), (B) three-fold axis channel ($Mg^{2+}$) and (C) four-fold axis channel ($Fe^{2+}/Fe^{3+}$).

### Supplementary Tables

| EMDB_id | Organism/Virus | Sample | Microscope | Filter | Resolution (Å) | Detector | Particles (N) | Reference | PDB_id |
|---|---|---|---|---|---|---|---|---|---|
| 9914 | *Mus musculus* | apoferritin | FEI TALOS ARCTICA | GIF | 2.0 | GATAN K2 SUMMIT | 323292 | [6] | n/a |
| 8743 | *Oryctolagus cuniculus* | aldolase | FEI TALOS ARCTICA | none | 2.6 | GATAN K2 SUMMIT | 83910 | [7] | 5vy5 |
| 6840 | *Escherichia coli* | beta-galactosidase with PETG | CRYOARM 200 | Omega | 2.6 | GATAN K2 IS | 93975 | n/a | n/a |
| **10205** | ***Mus musculus*** | **apoferritin** | **FEI GLACIOS** | **none** | **2.7** | **FEI FALCON III** | **95733** | **n/a** | **6sht** |
| 407 | *Homo sapiens* | methemoglobin | FEI TALOS ARCTICA | none | 2.8 | GATAN K2 SUMMIT | 24308 | [8] | 6nbc, 4n7p |
| 9671 | Ao-associated virus 2 | Virus, Ao-associated virus 2 | FEI TECNAI ARCTICA | n/a | 2.8 | FEI FALCON II | 14434 | [9] | 6ih9 |
| 4977 | Suppressed | Suppressed | FEI TALOS ARCTICA | n/a | 2.8 | FEI FALCON II | 56911 | n/a | 6rpk |
| 9672 | Ao-associated virus 2 | Virus, AAV2 with AAVR | FEI TECNAI ARCTICA | n/a | 2.8 | FEI FALCON II | 16820 | [9] | 6ihb |
| 9608 | Seneca valley virus | Virus, Seneca valley virus | FEI TECNAI ARCTICA | n/a | 2.8 | FEI FALCON II | 9167 | [10] | 6adm |
| 9612 | Seneca valley virus | Virus, Seneca valley virus | FEI TECNAI ARCTICA | n/a | 2.8 | FEI FALCON II | 13010 | [10] | 6ads |
| 406 | *Equus caballus* | Alcohol dehydrogenase | FEI TALOS ARCTICA | none | 2.9 | GATAN K2 SUMMIT | 11672 | [8] | 6nbb, 2jhf |
| 9706 | *Macrobrachium rosenbergii* nodavirus | Virus, nodavirus | FEI TALOS ARCTICA | n/a | 2.9 | FEI FALCON III | 19049 | n/a | 6jjc |
| 9798 | *Sulfolobus solfataricus* | ketol-acid reductoisomerase | FEI TALOS ARCTICA | n/a | 2.9 | FEI FALCON III | 34743 | [11] | 6jcv |
| 20133 | *Yersinia pestis* | Lon protease | FEI TALOS ARCTICA | n/a | 3.0 | GATAN K2 SUMMIT | 118143 | n/a | 6on2 |
| 552 | *Homo sapiens* | m-AAA protease AFG3L2 | FEI TECNAI ARCTICA | n/a | 3.0 | GATAN K2 SUMMIT | 1129437 | [12] | 6nyy, 6az0 |
| 4901 | Suppressed | Suppressed | FEI TALOS ARCTICA | n/a | 3.0 | FEI FALCON III | 43239 | n/a | 6rja |

Table S1. Cryo-EM reconstructions resolved at 200 kV < 3.0 Å. The data set described in this manuscript is highlighted in blue.

| EMDB_id | Organism | Microscope | kV | Resolution (Å) | Detector | Particles (N) | References | PDB_id |
|---|---|---|---|---|---|---|---|---|
| 9865 | *Mus musculus* | CRYOARM 300 | 300 | 1.5 | GATAN K2 SUMMIT | 120295 | n/a | n/a |
| 9599 | *Mus musculus* | FEI TITAN KRIOS | 300 | 1.6 | FEI FALCON III | 147000 | [6] | |
| 144 | *Escherichia coli* | FEI TITAN KRIOS | 300 | 1.7 | GATAN K2 SUMMIT | 426450 | [13] | n/a |
| 20026 | *Homo sapiens* | FEI TITAN KRIOS | 300 | 1.8 | GATAN K2 SUMMIT | 70648 | n/a | n/a |
| 10101 | *mus musculus* | FEI TITAN KRIOS | 300 | 1.8 | FEI FALCON III | 441902 | n/a | 6s61 |
| 9890 | *Homo sapiens* | CRYOARM 300 | 300 | 1.9 | GATAN K2 SUMMIT | 36855 | [14] | n/a |
| *9914 | *Mus musculus* | FEI TALOS ARCTICA | 200 | 2.0 | GATAN K2 SUMMIT | 323292 | [6] | n/a |
| 4905 | *Equus caballus* | FEI TITAN KRIOS | 300 | 2.1 | FEI FALCON III | 41202 | [15] | 6rjh, 4v1w |
| 4213 | *Homo sapiens* | FEI TITAN KRIOS | 300 | 2.1 | FEI FALCON III | 54006 | n/a | n/a |
| 263 | *Equus caballus* | FEI TITAN KRIOS | 300 | 2.2 | GATAN K2 SUMMIT | 65131 | [13] | n/a |
| 20027 | *Homo sapiens* | FEI TITAN KRIOS | 300 | 2.3 | GATAN K2 SUMMIT | 9600 | n/a | n/a |
| 3853 | *Homo sapiens* | FEI TITAN KRIOS | 300 | 2.5 | FEI FALCON III | 37000 | n/a | n/a |
| 20228 | *Homo sapiens* | FEI TITAN KRIOS | 300 | 2.5 | GATAN K2 SUMMIT | 304638 | [16] | 2ffx |
| 4701 | *Mus musculus* | JEOL 3200FSC | 300 | 2.7 | GATAN K2 SUMMIT | 184777 | n/a | n/a |
| **10205** | ***Mus musculus*** | **FEI GLACIOS** | **200** | **2.7** | **FEI FALCON III** | **95733** | **n/a** | **6sht** |
| 4698 | *Mus musculus* | JEOL 3200FSC | 300 | 2.8 | GATAN K2 SUMMIT | 190785 | n/a | n/a |
| 20227 | *Homo sapiens* | FEI TITAN KRIOS | 300 | 2.9 | GATAN K2 SUMMIT | 96631 | [16] | 2ffx |
| 20155 | *Equus caballus* | FEI TITAN KRIOS | 300 | 2.9 | DIRECT ELECTRON DE-64 (8k x 8k) | 129140 | n/a | n/a |
| 6800 | *Homo sapiens* | FEI TITAN KRIOS | 300 | 2.9 | FEI FALCON II | 32585 | [17] | n/a |
| 20229 | *Homo sapiens* | FEI TITAN KRIOS | 300 | 2.9 | GATAN K2 SUMMIT | 73314 | [16] | 2ffx |
| 20225 | *Homo sapiens* | FEI TITAN KRIOS | 300 | 2.9 | GATAN K2 SUMMIT | 72521 | [16] | 2ffx |
| 8428 | *Equus caballus* | FEI POLARA 300 | 300 | 3.0 | GATAN K2 SUMMIT | 4300 | [18] | n/a |
| 6802 | *Homo sapiens* | FEI TITAN KRIOS | 300 | 3.0 | FEI FALCON II | 63015 | [17] | n/a |

*reconstruction resolved at 200 kV, an energy filter was applied with a K2 direct electron detector.

Table S2. Reconstructions of apoferritin resolved at resolutions < 3.0 Å. The data set described in this manuscript is highlighted in blue.
